# Adeno-Associated Virus Technologies and Methods for Targeted Neuronal Manipulation

**DOI:** 10.1101/759936

**Authors:** Leila Haery, Benjamin E. Deverman, Katherine Matho, Ali Cetin, Kenton Woodard, Connie Cepko, Karen I. Guerin, Meghan A. Rego, Ina Ersing, Susanna M. Bachle, Joanne Kamens, Melina Fan

**Affiliations:** Addgene, Watertown MA 02472, USA; Stanley Center for Psychiatric Research, Broad Institute of MIT and Harvard, Cambridge, MA 02142, USA; Cold Spring Harbor Laboratory, Cold Spring Harbor, NY 11724, USA; Allen Institute for Brain Science, Seattle, WA 98109, USA; Penn Vector Core, Gene Therapy Program, University of Pennsylvania, Perelman School of Medicine, Philadelphia, PA 19104, USA; Departments of Genetics and Ophthalmology, Harvard Medical School, Howard Hughes Medical Institute, Boston, MA 02115, USA

## Abstract

Cell-type-specific expression of molecular tools and sensors is critical to construct circuit diagrams and to investigate the activity and function of neurons within the nervous system. Strategies for targeted manipulation include combinations of classical genetic tools such as Cre/loxP and Flp/FRT, use of cis-regulatory elements, targeted knock-in transgenic mice, and gene delivery by AAV and other viral vectors. The combination of these complex technologies with the goal of precise neuronal targeting is a challenge in the lab. This report will discuss the theoretical and practical aspects of combining current technologies and establish best practices for achieving targeted manipulation of specific cell types. Novel applications and tools, as well as areas for development, will be envisioned and discussed.

## Introduction

Understanding neural networks as they relate to development, behavior, and learning is a critical objective of neuroscience. These questions can be addressed, in part, by understanding the role of specific neural cells and brain regions, as well as the impact of individual molecules in these circuits. The successful execution of these neurobiology studies requires methods that are highly targetable, efficient, and precise. In this regard, recombinant adeno-associated viral vectors (herein referred to as AAV) are powerful tools that can be used both to target and manipulate specific neuronal subtypes (defined based on gene expression, location, and connectivity) and non-neuronal cell types within the nervous system.

Scientists using AAV for gene transfer and/or neuronal targeting must consider various questions about experimental design, including (1) how to best deliver/administer AAV (Fig. 1A), (2) which AAV serotype to use (Fig. 1B), and (3) how to drive gene expression with gene regulatory elements (both within the AAV genome and the host animal or cell line) (Fig. 1C-F). These and many other factors can affect how efficiently cells of interest are targeted by AAV. Further, experimental parameters such as AAV titer and dosage can impact AAV efficiency, and these details are often omitted from experimental methods in the literature and can be expensive and timely to determine empirically for each study. Overall, designing an experiment with AAV is multifaceted and suboptimal experimental design can drastically reduce the quality of results. In this report, we will discuss practical aspects of using AAV and considerations for designing experiments.

**Figure 1.**
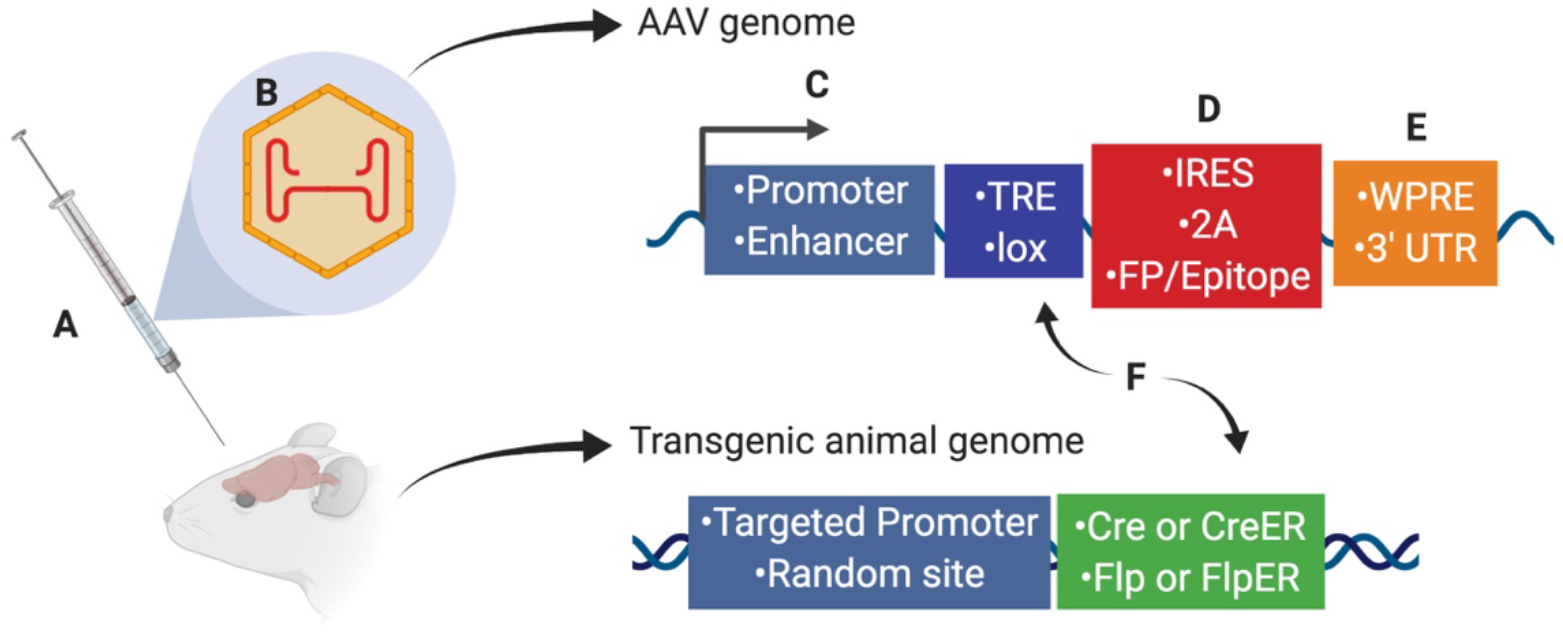
Several aspects of experimental design affect neuronal targeting and manipulation including (A) viral delivery method, (B) composition of viral capsid proteins, (C) promoters and/or enhancers driving transgene expression, (D) IRES or 2A elements for multicistronic expression coupled with fluorescent proteins (FP) or protein epitopes, (E) post-translational regulatory elements such as WPRE or 3’ UTR, and (F) Recombinase (Cre, CreER, Flp, or FlpER) expression from transgenic driver lines (inserted genomically via targeted or random integration) and ligand-dependent or recombinase-dependent expression elements such as TRE or lox sites, respectively. Abbreviations: TRE, tetracycline-response element; lox, LoxP sequence; IRES, internal ribosomal entry site; 2A, 2A sequence for self-cleavage; FP, fluorescent protein; WPRE, woodchuck hepatitis virus posttranscriptional regulatory element; 3’ UTR, 3’ untranslated sequence.

## Selecting the Route of Administration and Capsid

AAV tropism, as dictated by AAV capsid proteins, is an important factor affecting transduction efficiency and specificity across cell types. Since the mechanism of AAV transduction is through interaction of the AAV capsid with cell surface proteins and glycans, protein composition of the capsid (i.e., the AAV serotype) and the cell surface (i.e., based on cell type) determines transduction efficiency. Consequently, serotype and route of delivery should be carefully considered when designing experiments (Fig 1A, B). For an overview of the primary receptors for AAV serotypes, see (Schultz and Chamberlain, 2008).

### Direct intraparenchymal delivery

When injected directly into the brain, many of the naturally-occurring AAV capsids, which share homology ranging from 65-99% (Drouin and Agbandje-McKenna, 2013), have distinct but significantly overlapping tropisms and distribution characteristics. AAV1, AAV2, AAV5, AAV8, AAV9 and the engineered variant AAV-DJ are commonly used to target local populations of neurons after direct injections (Table 1). There are important differences in how far different capsid variants spread from the injection site - AAV2 and AAV-DJ diffusion is more confined and, therefore, these capsids are often chosen for applications that require precise targeting. While expression from AAV2 is mostly neuronal, several serotypes, including AAV1, AAV5, AAV8 and AAV9, also transduce astrocytes and oligodendrocytes.

**Table 1:** AAV administration routes for neuroscience

|  | Administration Route Expression |  |  |
| --- | --- | --- | --- |
|  | Direct | Intravenous | Delivery into the CSF (IT/ICV/CM) |
| Advantages | Regional expression<br>High levels of expression achievable (high MOI)<br>Requires small volumes of virus<br>Reduced off-target effects | CNS or PNS-wide transduction<br>Quick, noninvasive<br>Does not require surgical expertise<br>Lower more uniform expression<br>Sparse labeling is possible | IT injection can be used to target spinal motor neurons and dorsal root ganglia<br>Neonatal ICV injections can provide widespread gene delivery to the CNS<br>May allow CNS expression in the presence of neutralizing antibodies (Gray et al., 2013) |
| Disadvantages | Requires invasive surgery<br>Damage to the targeted area<br>Challenging in certain deep brain structures<br>Transduction gradient from injection site | Higher dose and volume of virus required<br>Greater risk of immune response<br>Off-target effects may confound experiment | Expression is not confined to the CNS<br>Requires moderately large volumes of virus<br>Expression is not as uniform as it is after systemic delivery |
| Expression considerations | High levels of expression may be important for opsin expression<br>High level expression makes cell-type specific transgene expression using regulatory elements more challenging | Moderate expression provided by IV AAV-PHP.B/eB may be preferable for GCaMP6 expression (no nuclear expression observed, see (Hillier et al., 2017)) | Expression is higher around CSF spaces and the brain/SC surface |
| Capsids | AAV2 – confined spread, mostly neuronal<br>AAV-DJ – Confined spread, higher expression (vs AAV2)<br>AAV1, 5 and 8 – widespread, moderate expression, neurons and glia<br>AAV2-Retro – widespread distribution, enhanced axonal uptake and retrograde expression<br>AAV1 – paired with Cre exhibits trans-synaptic (anterograde) transduction | AAV9 and rh.10 – efficient neonatal CNS transduction<br>AAV-BR1 – brain endothelial cell specific<br>AAV-PHP.B – enhanced neuron and glial transduction after adult IV injection in mice<br>AAV-PHP.eB – further evolved AAV-PHP.B variant with improved neuronal transduction<br>AAV-PHP.S – evolved capsid with improved transduction of peripheral nerves and heart | AAV7, AAV9 and Rh.10 are the most widely tested serotypes for delivery into the CSF (Borel et al., 2016; Federici et al., 2012; Gurda et al., 2016; Samaranch et al., 2013)<br>AAV4 enables transduction of ependymal cells (Liu et al., 2005)<br>AAV SCH9 and AAV4.18 enable SVZ progenitor cell transduction (Murlidharan et al., 2015; Ojala et al., 2018) |

### Systemic delivery

Several natural AAV capsids cross the blood brain barrier (BBB). In contrast to direct injections, intravenous injections of AAV can provide central nervous system (CNS)-wide gene delivery. This activity is present across several species and is most pronounced when AAV is administered to the neonate. Neonatal injections of AAV9 and rh.10 have been used to transduce neurons broadly across the CNS. However, when delivered at later developmental stages including in the adult, transduction is more limited and primarily restricted to endothelial cells and astrocytes, with transduction occuring in 1-2% of neurons in the forebrain (Deverman et al., 2016; Dufour et al., 2014; Foust et al., 2009). In this context, engineered AAV capsids have provided new and dramatically more efficient options for widespread gene delivery to the CNS. The first of these vectors, AAV-PHP.B, enabled researchers to deliver genes to more than 50% of neurons and astrocytes across numerous brain regions with a single noninvasive injection (Deverman et al., 2016). Achieving this efficiency requires relatively high viral doses (~1×10^14^ vector genomes/kg), thus requiring large volumes of high titer virus. A further-evolved AAV-PHP.B variant, AAV-PHP.eB, addresses this issue and can achieve >50% transduction of most neuron and astrocyte populations even with a 20-fold reduction in dose (Chan et al., 2017). While the activity of the PHP capsids is not universally observed across all species, the receptor engaged by the PHP capsids during AAV transduction has been identified and can be used to predict permissivities of cell or tissue types to these engineered capsids (Huang et al., n.d.). In addition, the same group has developed an additional AAV variant (the PHP.S variant) that can efficiently transduce dorsal root ganglia and other peripheral neuron populations following systemic administration, which should enable the study of these otherwise difficult to target peripheral neuron populations (Chan et al., 2017).

### CSF delivery

A third option for gene delivery to the CNS is to inject vectors into the cerebral spinal fluid (CSF). Several access points can be used: the lateral ventricle (intracerebroventricular, ICV), the cisterna magna (CM), subpial (Miyanohara et al., 2016) or the intrathecal (IT) space along the spinal cord. When performed in neonates, ICV AAV administration can provide widespread gene delivery. In the adult, ICV and CM injections result in gene delivery in multiple brain regions, however, the expression is not uniform across all brain regions and superficial structures are preferentially targeted. Beyond neurons, ICV injections also provide access to periventricular cell populations. For example, after ICV injection, AAV4 can be used to transduce the ependymal cells (Liu et al., 2005), and two engineered AAV capsids, SCH9 and AAV4.18, enable transduction of subventricular zone neural progenitors (Murlidharan et al., 2015; Ojala et al., 2018). IT injection can be used to deliver genes to spinal cord motor neurons and dorsal root ganglions (Federici et al., 2012; Foust et al., 2013; Schuster et al., 2014).

### Retrograde and anterograde transport for circuit studies

AAV vectors are also commonly used as part of circuit studies. Numerous natural AAV serotypes exhibit retrograde trafficking activity from their uptake at axon terminals. However, retrograde transduction with natural serotypes such as AAV1, AAV2, AAV6 and AAV9 requires high vector doses due to the relative inefficiency of this transduction mechanism. More recently Tervo et al. (Tervo et al., 2016) and Davidsson et al. (Davidsson et al., n.d.) have created modified capsids AAV2-Retro and AAV MNM008, respectively, that provide efficient transduction of neurons that send axon projections into the injection site. Transduction efficiencies of both capsids are shown to circuit-dependent, and thus capsids should be validated for circuits of interest when planning experiments.

In summary, a consensus of opinion has not been reached regarding the best serotype for each cell type, brain region or application. Choosing the optimal serotype requires reviewing the literature most relevant to the planned experiment and performing pilot testing for new or at least for challenging applications. As new engineered capsids with unique features continue to be developed, the available options will become more numerous and more powerful (Chan et al., 2017; Davidsson et al., n.d.; Deverman et al., 2016; Ojala et al., 2018; Tervo et al., 2016)

## Controlling Gene Expression with Regulatory Elements

Cre and Flp recombinase-dependent expression elements within AAV vectors remains the go-to system for restricting transgene expression to genetically defined cell types in model organisms. However, few Cre or Flp transgenic lines have been developed in other mammalian species. Furthermore, breeding multiple transgenic lines to generate the desired offspring can be time consuming and expensive. Therefore, there is significant interest in developing the means to achieve similar expression specificity in nontransgenic animals using flexible vector-based approaches that will translate across species.

Cis regulatory elements can be used to control transgene expression from AAV genomes. These elements include promoters and enhancers (Fig. 1C), as well as introns, micro-RNA recognition sequences, and internal ribosome entry sites (IRES) (Fig. 1D) that can be used to tailor RNA processing, stability, and translation to the experimental needs. Here we will discuss how these regulatory elements can be used to restrict AAV-mediated gene expression.

### Enhancers and promoters

Enhancer and promoters (hereafter referred to as promoters for simplicity) can generally be divided into two classes: general/ubiquitous and cell type-specific. Typically, ubiquitous promoters provide high-level, long-term expression in most cell types, though some, such as the viral CMV promoter, have been shown to exhibit silencing in specific tissues over time (Klein et al., 1998; McCown et al., 1996; Paterna et al., 2000; Tenenbaum et al., 2004) (Table 2). High expression levels are not optimal for every application and alternative regulatory elements, such as those from the mammalian MeCP2 or PGK genes (Table 3) may be suitable for experiments where high-level viral enhancer driven expression is not desired. General recommendations for expression levels of various types of transgenes is summarized in Table 4.

**Table 2:**
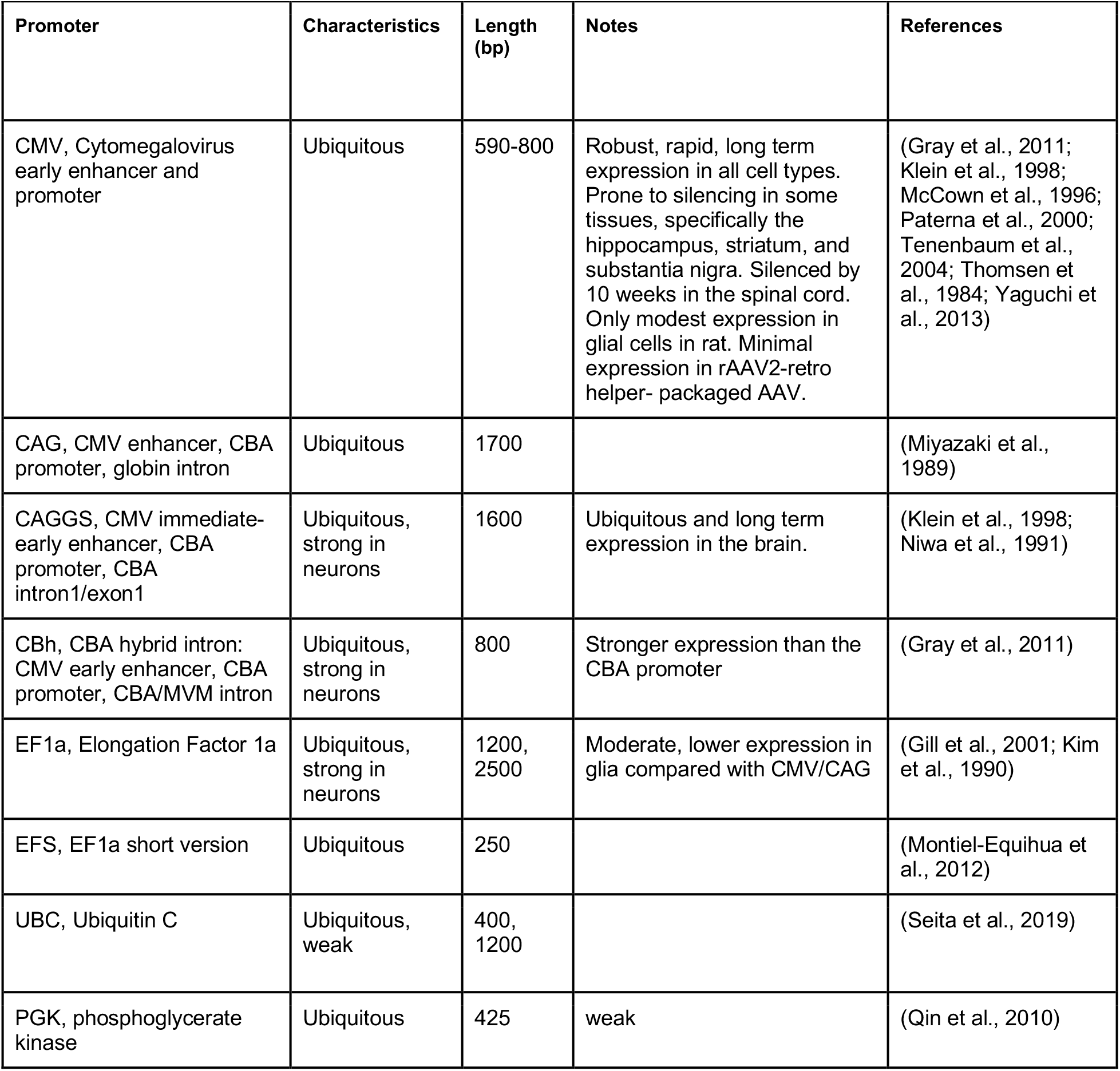
Ubiquitous enhancers and promoters

**Table 3:**
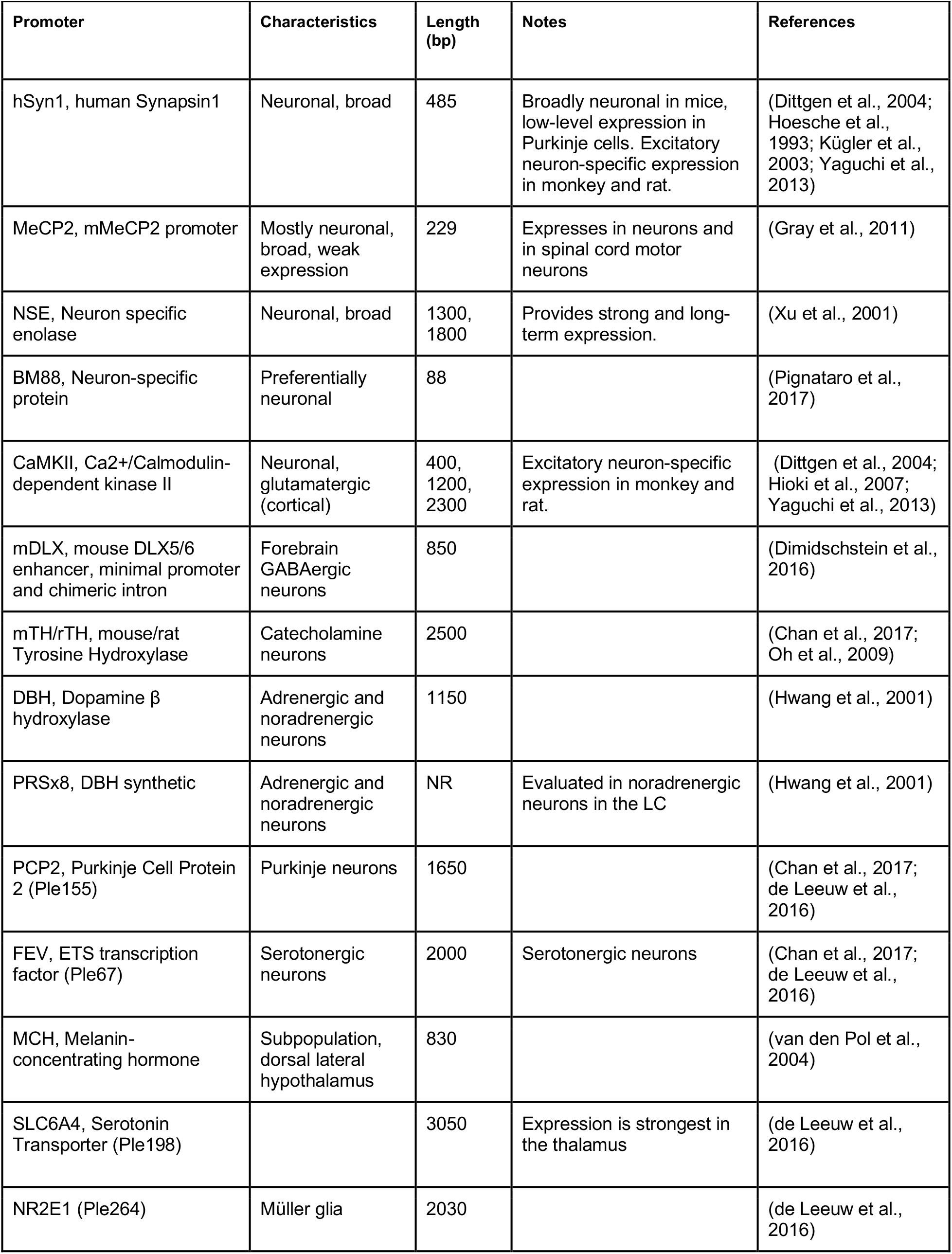
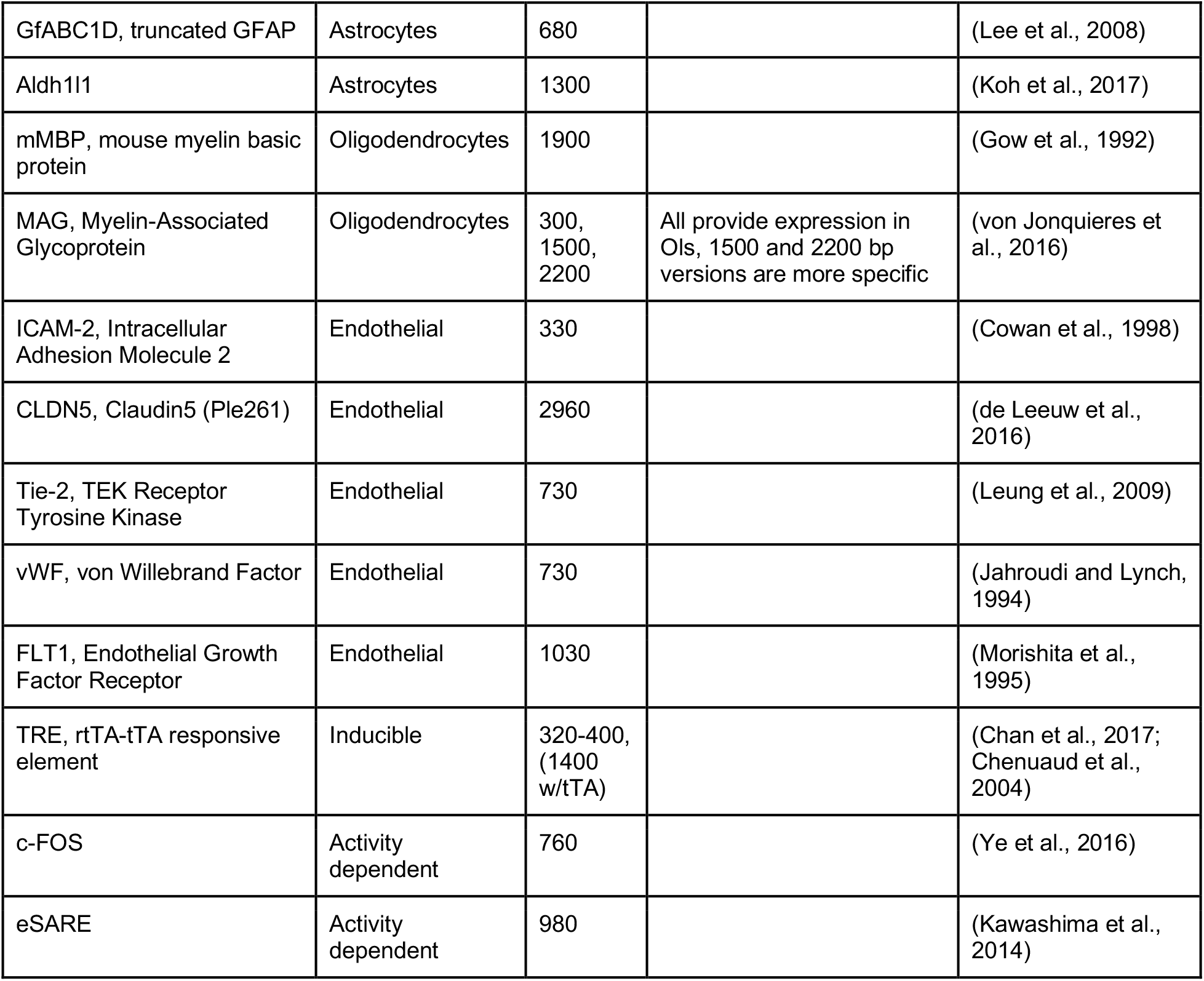
Cell type-specific promoters

**Table 4:** General expression considerations for specific transgenes and applications

| Transgene | Application | Optimal expression level |
| --- | --- | --- |
| Opsins | Optogenetics | High |
| DREADDs | Chemogenetics | Low for optimal specificity |
| Ca<sup>2+</sup> sensors | Activity monitoring | Moderate |
| Voltage sensors | Activity monitoring | Moderate-high |
| dLight1 | Dopamine indicator | Not yet known |
| Fluorescent reporters (XFPs) | Expression reporters, protein tagging | Moderate |
| Luciferase/AkaLuc | Expression reporter | Low-High |
| Cre/FlpO/Dre/KD/B3 Recombinases | Intersectional expression/<br>Circuit studies | Low |
| CRISPR-Cas9 | Gene editing | Moderate-High |
| TVA and rabies G | Circuit studies/retrograde tracing/TRIO/cTRIO | TVA (low)<br>Rabies G (Moderate) |

Viruses evolved under constraints that drove the selection of short, strong promoters. In contrast, the regulatory elements that control mammalian gene transcription are often distributed over thousands to hundreds of thousands of bases. Due to the limited packaging capacity of the AAV genome, identifying AAV-compatible promoters has been challenging and the development of shortened promoters is an area of active study (de Leeuw et al., 2016; Nathanson, 2009). A list of cell type-specific promoters compatible with AAV vectors is provided in Table 3.

### Multicistronic vectors

Although AAV vectors have a limited packaging capacity, it is possible to express multiple short transgenes from a single vector using one of several approaches: (1) separate translational units where each cDNA is controlled by separate 5’ and 3’ regulatory elements, (2) using IRES sequences to insert two separate translational units into a single mRNA, or (3) the use of viral 2A sequences to generate separate proteins from the same translational unit (Fig. 1D). Table 5 highlights considerations for choosing between the use of IRES sequences and 2A “selfcleaving” peptides.

**Table 5:** Bicistronic Expression Options

|  | Bicistronic Expression Elements |  |
| --- | --- | --- |
|  | 2A | IRES |
| Advantages | Requires minimal sequence space<br>Results in similar expression of both proteins<br>Can be used to express >2 proteins | Protein products are unmodified |
| Disadvantages | May reduce expression of both proteins<br>Adds peptide sequence to the C-terminus of the first protein and a proline to the N-terminus of the second protein | May not provide equivalent expression of both transgenes<br>IRES sequences are >500 bp |

One important consideration when evaluating expression strategies and determining specificity is that the expression levels required for reporter detection may not match what is necessary for the activity of an opsin, DREADD or recombinase. For example, fluorescent proteins are commonly used to evaluate gene regulatory elements and vector design. However, fluorescent reporter assays may give the false impression of specificity if high levels of expression are seen in target cell types and low-level expression goes undetected in off-target populations. If these same regulatory elements are then used to drive DREADDs or Cre, which can mediate their effects at low expression levels, then the specificity may appear reduced. If this goes unexamined, then the interpretation of experimental results could be compromised. Moreover, though they are commonly used, fluorescent proteins are not necessarily inert and can lead to immune responses in larger animals, and over-expression related toxicities in mice.

### Post-translational regulatory elements

Transgene expression can also be controlled post-transcriptionally through the use of elements impacting RNA splicing, nuclear export, stability, and translation into proteins (Fig. 1E). Inclusion of an intron can have positive impacts on expression levels. Introns have also been combined creatively with recombinase sites and partially inverted transgenes to achieve tight intersectional control of transgene expression (Fenno et al., 2017, 2014). Many recombinant AAV genomes also include a woodchuck hepatitis virus posttranscriptional regulatory element (WPRE), which can dramatically enhance expression. For several examples of how inclusion of a WPRE affects expression from AAV vectors please see (de Leeuw et al., 2016).

Complementary miRNA target sites (TS) are frequently engineered into the 3’ untranslated region (3’ UTR) of AAV genomes to mitigate off-target transgene expression. These sequences are complementary to miRNAs expressed within off-target cell types but not within the target population. miRNA binding to the perfectly complementary miRNA TS results in degradation of the RNA. Inclusion of multiple copies of the short miRNA TS sequences can dramatically lower off-target transgene expression. For example, by incorporating three copies each of miR-1 and miR-122, which are specifically expressed in muscle and liver, respectively, Xie et al. reduced transgene expression from intravenously administered AAV9 in muscle, heart, and liver, while maintaining brain expression (Xie et al., 2011). miRNAs that enhance the restriction of lentiviral mediated gene expression to GABAergic neurons have also been identified (Keaveney et al., 2018; Xie et al., 2011). Given their short lengths, miRNA TS can be multiplexed within the same genome, making them attractive elements for reducing expression outside of the cell type of interest.

### Knock-in driver Cre and Flp mouse lines

The mammalian central nervous system contains extremely diverse types of neurons. These neuronal cell types can be distinguished by their intrinsic gene expression profiles, which are differentially regulated throughout development. Binary expression systems of drivers and reporters can be used to drive transgene expression in genetically-defined cells (reviewed in (Huang et al., 2014)).

There are two general strategies for using binary expression systems to access genetically-defined cell types: transactivation-based systems (based on tet-response elements, TRE) and recombination-based systems (based on lox-site recombination, lox) (described in (Taniguchi, 2014)) (Fig. 1F). Gene targeting techniques can be used to insert *Cre* or *Flp* site-specific recombinases in the mouse genome. These knock-in driver mouse lines express Cre or Flp under the activity of a target gene’s endogenous promoter. Thus, Cre and Flp driver mouse lines constitute a genetic switch to turn on a recombinase-dependent reporter or effector. Since the recombinase is only expressed in cells defined by the target gene’s endogenous promoter activity, this system allows labeling and manipulation of neurons defined by the targeted gene’s expression pattern.

The reporter or effector whose expression is dependent on the driver’s activity is introduced in vivo by either crossing the driver to a transgenic reporter mouse line, using a viral vector, or electroporating the DNA construct into the cells. For example, with the widely used Cre driver, the conditional reporter expression depends on Cre-dependent recombination of specific lox sites.

#### Targeting at random versus targeting to a specific gene’s locus

Knock-in Cre and Flp recombinases can be inserted in the genome either randomly or at a particular gene locus (Fig. 1F). Conventional transgenic and BAC transgenic approaches are targeted at random into the genome. However, knock-in mouse lines targeted to a specific locus by homologous recombination have the advantage that expression of the inserted recombinase will recapitulate the expression pattern within cells of the endogenous gene of interest. There are several advantages to using targeted knock-in Cre driver mouse lines (Table 6). Targeted knock-in Cre or Flp mouse lines can be inserted at the gene’s transcription/translation initiation site. When using this strategy, it should be noted that individuals can exhibit variable levels silencing. Typically, an optimal “non-silencing” male should be identified and used for genetic crosses. However, the offspring of this male may exhibit silencing and must be revalidated.

**Table 6:** Methods of delivering Cre for cell-type targeting, labeling, and manipulation.

|  | Method of Cre delivery |  |  |
| --- | --- | --- | --- |
|  | Targeted knock-in Cre mouse line | Transgenic Cre mouse line (not generated by homologous recombination) | Cre via AAV |
| **Advantages** | • Specific and reliable by genetic targeting to locus of interest (higher certainty that driver activity will reflect the endogenous expression of the gene of interest)<br>• Comprehensive with Cre mouse lines<br>• Sparse if using CreER by adjusting tamoxifen dose<br>• Can combine with viral strategies to achieve spatial control or very strong expression<br>• Lower Cre expression than AAV | • Cheap and easy to produce mouse lines<br>• Lower Cre expression than AAV | • Stable over time<br>• Spatial control: can restrict delivery to a particular region<br>• Can be delivered broadly by systemic (e.g., tail vein or retro-orbital) injection |
| **Disadvantages** | • Time-consuming and costly to produce and maintain mouse lines | • Does not necessarily recapitulate the endogenous gene's expression<br>• Genetic silencing in mouse lines can affect Cre expression<br>• Sexual dimorphism can arise that does not reflect the gene's native expression profile<br>• Transgenic animals can lose specificity over time | • Expression gradients from injection site(s)<br>• AAV vectors increase interleukin levels<br>• Cre toxicity if delivered at high dose<br>• AAV expression may affect the cell's DNA and RNA |

#### Temporal control with tamoxifen

To control the expression of a reporter in a subset of neurons that uniquely and transiently expresses a certain marker at a particular time point, tamoxifen-inducible recombinases (Feil et al., 2009), so-called CreER or FlpER, can be used as drivers. In CreER driver mice, activation of the expressed Cre recombinase requires administration of the estrogen receptor modulator drug tamoxifen to the animal, allowing for temporal control of the subpopulation.

The typical tamoxifen dose varies from 10-200 mg/kg, depending on the desired degree of recombination at the reporter allele. In effect, this controls the sparseness/density of the labeling of the targeted neuronal population. The timing for tamoxifen administration depends on the temporal characteristics of the promoter driving CreER in the specific population to be targeted. Importantly, the half-life of tamoxifen (approximately 48 hours) must be taken into consideration. Tamoxifen preparation is detailed in the published protocol in (Vaughan et al., 2015).

Tamoxifen can be administered in one of three ways, depending on the desired developmental time point for activating the CreER driver: oral gavage to the mother for embryonic induction; subcutaneous injection to offspring for early postnatal induction, or intraperitoneal injection in offspring for late postnatal and adult induction.

#### Best practices for induction of CreER or FlpER mouse lines

To generally improve the reliability of results obtained with CreER or FlpER mouse lines, tamoxifen dosage should be varied. If the level of recombination of the reporter achieved with a high dose of tamoxifen is low, administration of 4-hydroxytamoxifen (the native form of tamoxifen) can improve activation of CreER (Jahn et al., 2018). Tamoxifen needs to be metabolized in the liver to reach its active form so it can act on CreER. However, it can be preferred over 4-hydroxytamoxifen due to lower cost. Consider that the CreER or FlpER will be active for 48 hours following tamoxifen induction, which may impact induction during developmentally active time periods. Administration of tamoxifen by gavage to the pregnant mother to induce the pups at embryonic timepoints can lead to problems of miscarriage or poor mothering. When administering tamoxifen at embryonic timepoints, use of Swiss or CD1 compared to C57Bl6 mice can improve outcome in two ways: they produce larger litters and females are better mothers, which overall can improve pup survival.

#### Validating a knock-in driver mouse line

Knock-in Cre driver mouse lines are designed to control expression of reporter probes in genetically-defined cell-types. It is important to validate that the driver mouse line expressing a site-specific recombinase (e.g., Cre, CreER, Flp, FlpER) reflects the endogenous expression pattern of the gene targeted by the knock-in driver. Various approaches can be used separately or jointly to validate a knock-in driver mouse line: (1) Crossing the mice with a suitable reporter like Rosa26-CAG-LSL-td-tomato (Ai 14) or Rosa26-CAG-LSL-h2B-GFP and assessing brainwide expression, (2) Immunostaining of the target regions, (3) dual fluorescent in situ hybridization (dual fISH) with probe for reporter (e.g., RFP or GFP for Rosa26-CAG-LSL-td-tomato (Ai14) or Rosa26-CAG-LSL-h2B-GFP, respectively). Note that assessing Cre lines by crossing to a reporter line gives an integrated view of Cre activity over the lifetime of the animal. To assess Cre activity in the target cell population at the particular age of interest, a viral vector with a Cre-dependent reporter in a Cre line can be used. Similarly, for CreER lines tamoxifen induction should be performed at different developmental timepoints to assess temporal specificity.

In addition, leakiness of both mouse lines and viruses must be assessed prior to interpretation of experimental results that rely on the complete restriction of off-target Cre and/or transgene expression. Leakiness of a mouse line can be assessed when validating the mouse line, ensuring that expression of the site-specific recombinase or activator (i.e., Cre, CreER, Flp, FlpER or tTA) is consistent with the gene of interest in its native form. Crossing the driver to a reporter (e.g., fluorescent reporter) and performing dual fluorescent in situ hybridization (dual fISH), with probes for the fluorescent reporter and for the gene of interest allows one to check that endogenous expression of the gene and activity of the driver are present in the same cells. Using knock-in driver mouse lines instead of BAC driver mouse lines allows for targeted expression, with a higher likelihood of expression within cells expressing the endogenous gene of interest. Leakiness of the tamoxifen-inducible driver mouse line can be assessed by crossing the inducible driver CreER or FlpER to a Cre or Flp dependent fluorescent reporter, and checking for expression of the fluorescent protein without administering tamoxifen. Finally, leakiness of the Cre-or Flp-dependent or tTA-activated AAV can be checked by injecting the virus in a mouse that has been crossed to a fluorescent reporter mouse line for the driver/activator. If expression from the virus and from the mouse reporter line match, this indicates that the AAV is specific to the driver/activator.

## Viral Strategies for Targeting Defined Populations

Combining various experimental techniques can enable the precise targeting of specific neuronal populations of interest. The general advantage of combinatorial approaches is specificity of cell targeting can be improved with each additional technique. Using the natural expression patterns of AAV, neuronal populations can be specifically targeted based on their locations and connectivity. Here we will review several targeting strategies.

### Axon Terminals

Axon terminals can be targeted by allowing robust expression of the AAV payload through to the axons of neurons at the injection site. Then, terminals in specific areas can be targeted optogenetically by only delivering light to the region harboring the terminals of interest (Fig. 2a). This technique can be further restricted such that only axons of a particular cell type are targeted by using cell-type specific promoters to drive the AAV expression (Fig. 2b). Thus, specific axonal targeting can be achieved by coupling anterograde expression from a local injection with optogenetic stimulation at an axon terminal (Nassi et al., 2015).

**Figure 2.**
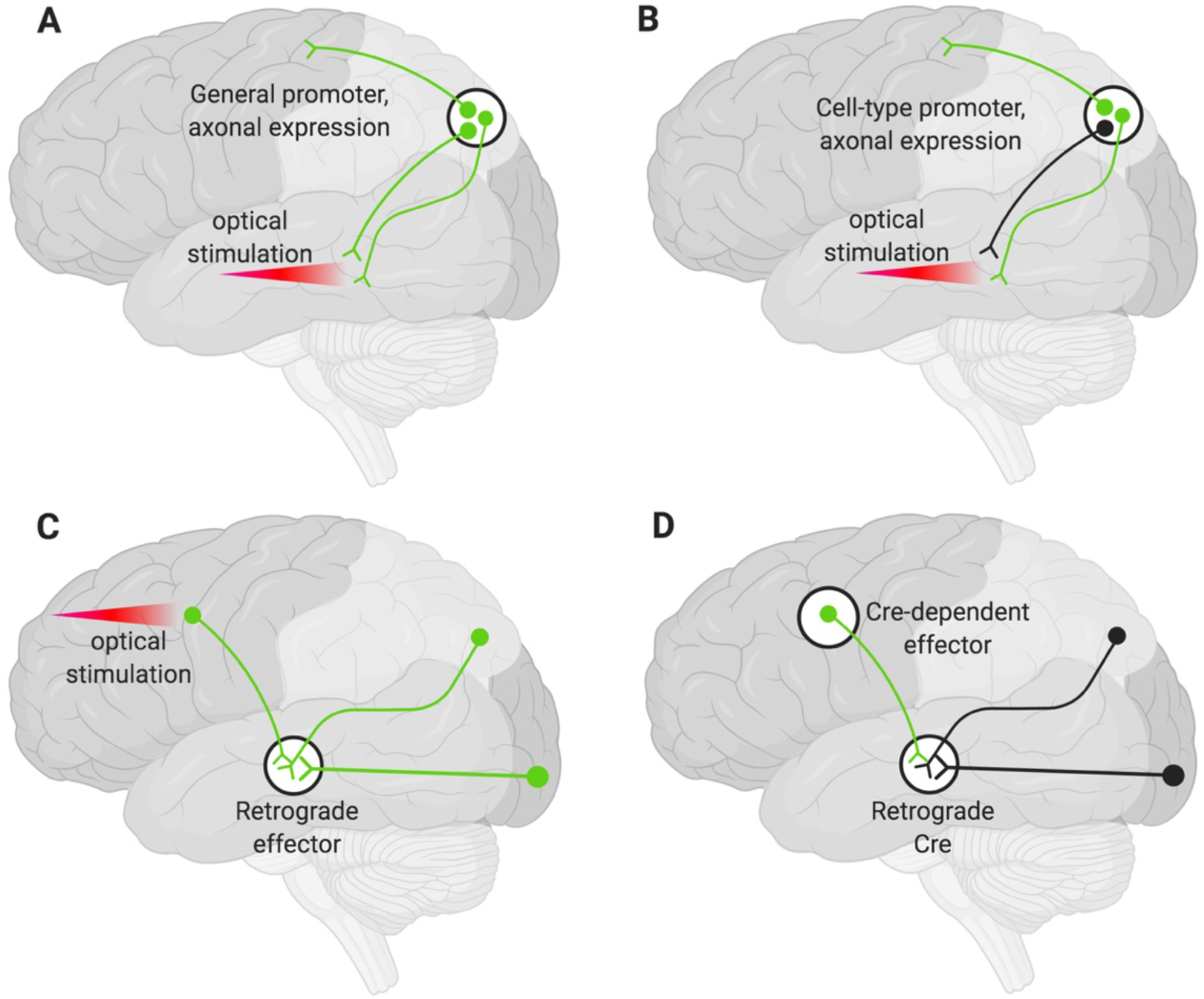
Various strategies for neuronal targeting using AAV. Delivery of neuronal effectors via AAV labels (green) axons and terminals with cell bodies at the injection site. (A) Effectors under a general promoter express in all transduced neurons with cell bodies at the injection site. Specific regions can be optically stimulated (red beam). (B) Effectors under cell-type promoters express only within a cell type. (C) Effectors delivered via a retrograde AAV express in all transduced neurons with axons that project into the injection site. Cell bodies in regions of interest can be optogenetically stimulated (red beam). (D) Delivery of a retrograde AAV expressing Cre recombinase (Retrograde Cre) to the projection site coupled with local delivery of a Cre-dependent effector limits expression to neurons within specific circuits.

### Projection Neuron Targeting

Targeting the cell bodies of projection neurons can be important for manipulating only those neurons that terminate at a particular site. To target these neurons, retrograde AAV can be delivered to projection sites and the cell bodies neurons terminating at that region will be targeted. This method alone will not give rise to pathway specificity. However, if used to deliver optogenetic tools, light can be delivered only to the region harboring the cell bodies of interest (Fig. 2c) (Nassi et al., 2015).

Finally, specific populations of projection neurons can be targeted by coupling local delivery of a Cre-dependent, inducible neuromodulator (e.g., a DREADD or opsin) with retrograde delivery of Cre to the site where the targeted projection neurons originate (Fig. 2d). Using this approach, a subset of projection neurons (Fig. 2d), rather than all the projection neurons (Fig. 2c), then express the Cre-dependent neuromodulator. This has the advantage that effectors (e.g., the DREADD ligand) can then be delivered systemically rather than locally and still only manipulate the subset of projection neurons that have been targeted (Nassi et al., 2015).

## Areas for Development

The arsenal of new sensors, actuators, recombinases, genes, RNA and base editing enzymes, and other genetically encoded tools for studying the nervous system is rapidly growing. AAV vectors remain the most versatile and powerful approach for delivering these tools to the CNS. Nevertheless, delivery challenges remain and efforts are ongoing to develop new vectors that address several key needs including (1) improved widespread CNS gene transfer via IV and ICV routes, (2) AAV vectors capable of more efficient anterograde transport, (3) vector solutions for delivering transgenes too large to fit in a single AAV virus, (4) capsids that specifically target defined neural cell types and neuronal subtypes, and (5) viral vectors that enable transduction of microglia.

Overall, consistency and repeatability of both existing and newly developed AAV tools can be improved by following best practices and guidelines. While powerful technologies are being developed, each has technical limits which need to be considered both when designing experiments and when interpreting results. To this end, improved platforms for sharing information, including technical guidelines and best practices, will serve the research community by enabling technologies to be used to their fullest capacities consistently across labs.

## Acknowledgements

We thank Gowan Tervo, Sarada Viswanathan, and John Dingus for helpful discussions. We are grateful for the generous support by The Kavli Foundation to hold the meeting that gave rise to this report. We also thank the Research Team and all other members of Addgene for advice and support during the preparation of this manuscript. Figures in this manuscript were created with Biorender.

